# Can public good producing subclones invade a population of non-producers?

**DOI:** 10.64898/2026.05.14.725277

**Authors:** Svyatoslav Tkachenko, Michael Hinczewski, Christopher D McFarland

## Abstract

Cancer progression is increasingly understood as an evolutionary process shaped not only by competition but also by cooperative interactions including those mediated through diffusible “public goods” (PGs). Classical evolutionary game theory predicts that PG-producing (altruistic) subclones cannot invade well-mixed populations of non-producers, creating a paradox given their observed emergence in tumors. Here, we resolve this contradiction by combining stochastic spatial simulations with an analytically tractable Moran model to study the invasion dynamics of PG-producing cells in structured populations. Starting from a single producer cell, we explicitly model stochastic PG secretion, diffusion, binding/unbinding, and cell proliferation across biologically relevant parameter ranges. We demonstrate that spatial structure fundamentally alters invasion dynamics, enabling PG producers to invade and establish even when production incurs a fitness cost. Both numerical and analytical approaches converge on a key unifying parameter, a characteristic length scale *δ*, that captures the combined effects of diffusivity, binding kinetics, and degradation. This length scale determines the spatial extent of PG availability and thus the selective advantage of producers. We identify distinct regimes: when PGs are localized (small *δ*), producers preferentially benefit and invasion is likely; when PGs are widely dispersed (large *δ*), benefits are shared and invasion approaches neutrality or is suppressed by costs. Our results highlight that invasion of cooperative traits is governed by spatially mediated resource localization rather than intrinsic fitness alone. This framework provides a mechanistic basis for understanding the emergence of cooperative subclones in tumors and suggests that modulating biophysical transport properties of signaling molecules could influence tumor evolution, metastasis, and therapeutic resistance.

## Introduction

Since the 1970s, cancer progression has been described as a Darwinian selection process driven by somatic cellular competition1–^5^. In recent years, however, we have learned that this process is not simply “survival of the fittest”: normal tissues are nicely-organized, fine-tuned orchestras whose coordinated melody is cunningly co-opted by tumorigenic cells^6–8^.

Tissue-level tumorigenic phenotypes are traditionally modeled using evolutionary game theory (EGT) instead of the traditional population genetics formalism, as natural selection no longer acts directly on the inherited cellular trait^9–13^. In these models, paracrine and juxtacrine signaling interactions between distinct tumor clones and stroma mediate clone co-existence and cooperation^9–15^. EGT modeling explains patterns of intratumor heterogeneity, proliferation, therapeutic targeting, and treatment resistance that cannot be explained by the traditional approach^7, 9, 16–20^. Still, EGT approaches often use well-mixed and infinite population approximations^21–23^, despite the fact that these interactions are known to be mediated by spatially-localized, soluble chemokines^6, 7, 11^. Moreover, EGT often relies on deterministic methods^23^ to understand how ‘established’ clones (i.e. clones that have avoided extinction via genetic drift) persist in a population.

How clones invade tumor in the first place and eventually establish persistent populations is understudied. In particular, the invasion of cells that produce an “altruistic” public good (PG)—*e*.*g*. chemokines that benefit the cell itself and competition alike—seems paradoxical in existing formalism: producer subclones should not be able to invade well-mixed populations^10, 13^. Indeed, the emergence of altruistic traits in general has intrigued evolutionary theorists for generations. Regardless of the apparent theoretical paradox, altruistic producer traits do arise in cancer^24–26^.

To explore this contradiction, we applied numerical and analytical methodologies to identify factors leading to invasion and establishment of a PG-producing subclone into a spatial tumor population. We consider stochastic effects in cell division, diffusion, binding and absorption of PG to ascertain the invasion probability of producers across a broad parameter space. Despite differences in the assumptions and approximations of our numerical and analytical models, we observe similar model behavior and define a characteristic length that determines invasion probability from the tissue dimensionality, and the diffusion, binding and unbinding constants of the PG. When spatial and stochastic properties of PGs are considered, the invasion of PG traits is possible for biologically-realistic parameters even when PG production confers a growth cost.

## Results

A minimal model of PG production and invasion into an evolving cellular population must include stochastic production, diffusion and binding of PG, alongside stochastic cell birth and death. We model these processes using both (i) numerical simulations that realistically captures all stochastic processes, and (ii) an analytical formalism that identifies powerful abstractions and offers general predictions for arbitrary dimensionality of tissue growth and fitness cost. Both models begin with a single PG producing cell within a sea of non-producers, and no other relevant mutations in the population (*i*.*e*. low mutation rate limit). We recently modeled the occurrence of many ecologically-interacting mutations in well-mixed populations and predicted dramatically different levels of genetic diversity and modes of drug resistance^27^; however, in this study we are investigating whether a *single* public good adaptation can emerge, if at all, in tumor populations.

### Spatial stochastic model of public good production and invasion

An invading cell begins at the center of the lattice (Fig. 1) and produces diffusible PG. Timescale analysis indicates that chemical diffusion and PG binding/unbinding processes are much faster than cellular division and death.

**Figure 1.**
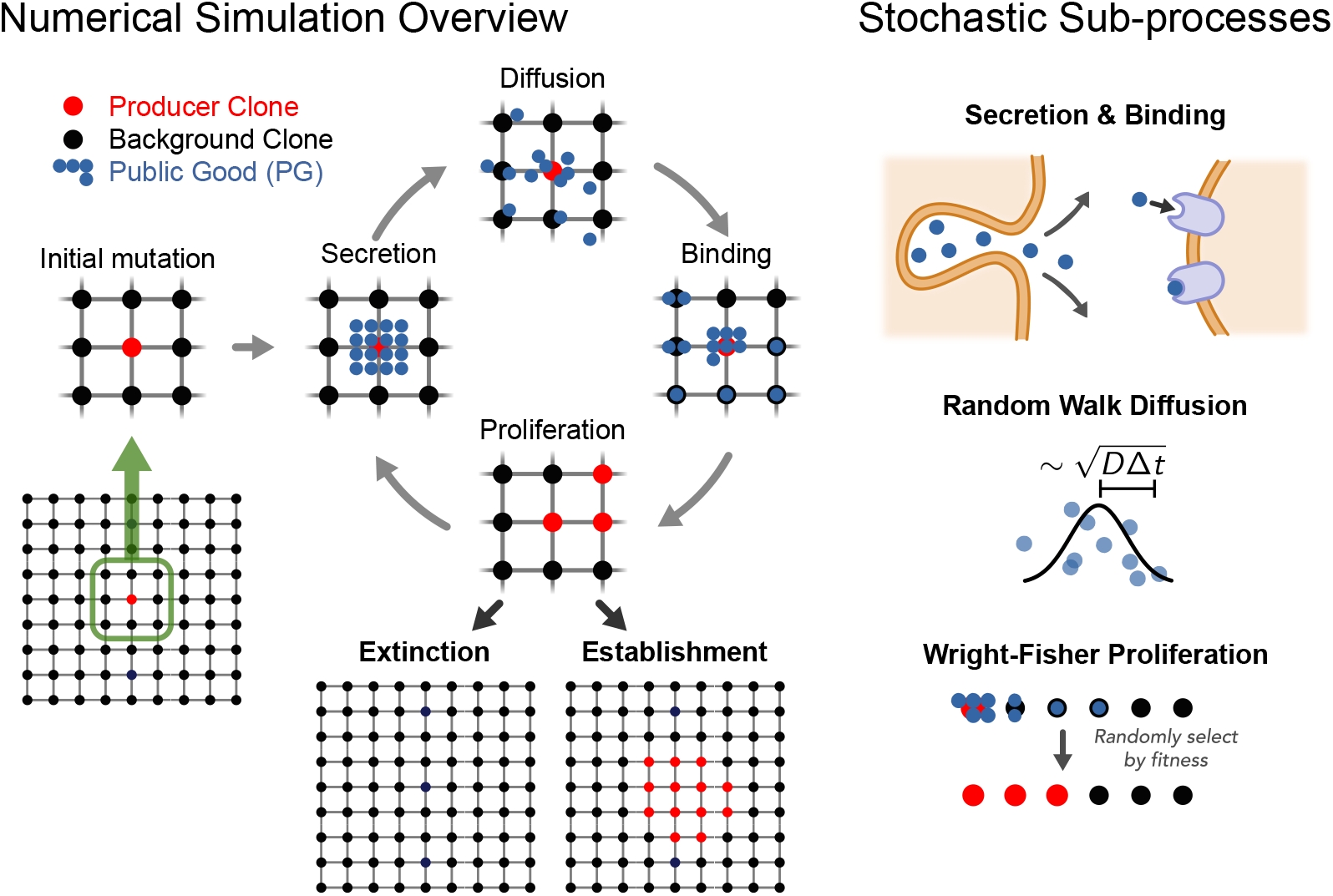
Overview of the numerical simulation. The initial mutation occurs in the middle of the lattice. The mutated cell secretes diffusible public good (PG) molecules. The molecules diffuse along lattice edges, stochastically binding to cells located at lattice nodes. Cells then proliferate according to the Wright-Fisher procedure with cell fitness determined by the number of PGs bound to each cell. The cycle is repeated until either public good producers go extinct or they occupy at least 5% of the nodes.

Hence, in our simulations PGs diffuse via a random walk on the lattice. Diffusing PGs may bind/unbind from cells on the lattice at each time step with association/dissociation rates *k*_on_ and *k*_off_. Association and dissociation are performed using a stochastic Gillespie algorithm, which is run at every diffusion step for the length of time it takes a PG to diffuse the length of the lattice edge.

Proliferation is modeled every 24 hours via a Wright-Fisher procedure. Offspring are selected in proportion to cell fitness *f*, which is a Hill function of bound PG:

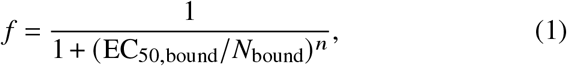

where EC_50,bound_ is the half-maximal effective concentration EC_50_ converted to the number of ligands providing that concentration; *N*_bound_ is the number of bound ligands; *n* is the Hill coefficient. This relationship between PG and fitness was proposed previously as it accurately models known signaling cascades^9^. To maintain spatial structure, all producer offspring are grouped together at the center of the lattice.

We assume that clones establish stable populations or “invade” (Methods) when the number of producers exceeds 5% of the total population. All simulations continue until a producer subclone invades or goes extinct (no invasion). We verified for a few parameter combinations that 5% and 50% thresholds gave identical results, indicating that a 5% population fraction of producers is a relevant threshold for allele establishment.

We then determined the probability of invasion for cells across all relevant parameters for five decades of parameter values reflecting experimentally measured value ranges^28^ (Table 1). The results are shown as heat maps of establishment probability, fixing one parameter and varying the other two, in the panels of Fig. 2. Unlike models assuming well-mixed populations, where the invasion of a single PG producer cell is impossible or exceedingly rare^13^, we find that invasion is possible—and probable— for many parameter combinations. Importantly, neither expanding population size nor complicated deme structure is needed in our system.

**Table 1.** Parameter values used in numerical simulations.

|  |  |  |  |  |  |  |  |  |  |  |
| --- | --- | --- | --- | --- | --- | --- | --- | --- | --- | --- |
| $D [\mu\text{m}^2/\text{s}]$ | 47 | 90 | 134 | 178 | 222 | 265 | 309 | 353 | 395 | 440 |
| $k_{\text{on}} [\text{M}^{-1}\text{s}^{-1}]$ | $10^3$ | $10^4$ | $10^5$ | $10^6$ | $10^7$ | $10^8$ | | | | |
| $k_{\text{off}} [\text{s}^{-1}]$ | $10^{-1}$ | $10^{-2}$ | $10^{-3}$ | $10^{-4}$ | $10^{-5}$ | $10^{-6}$ | | | | |

**Figure 2.**
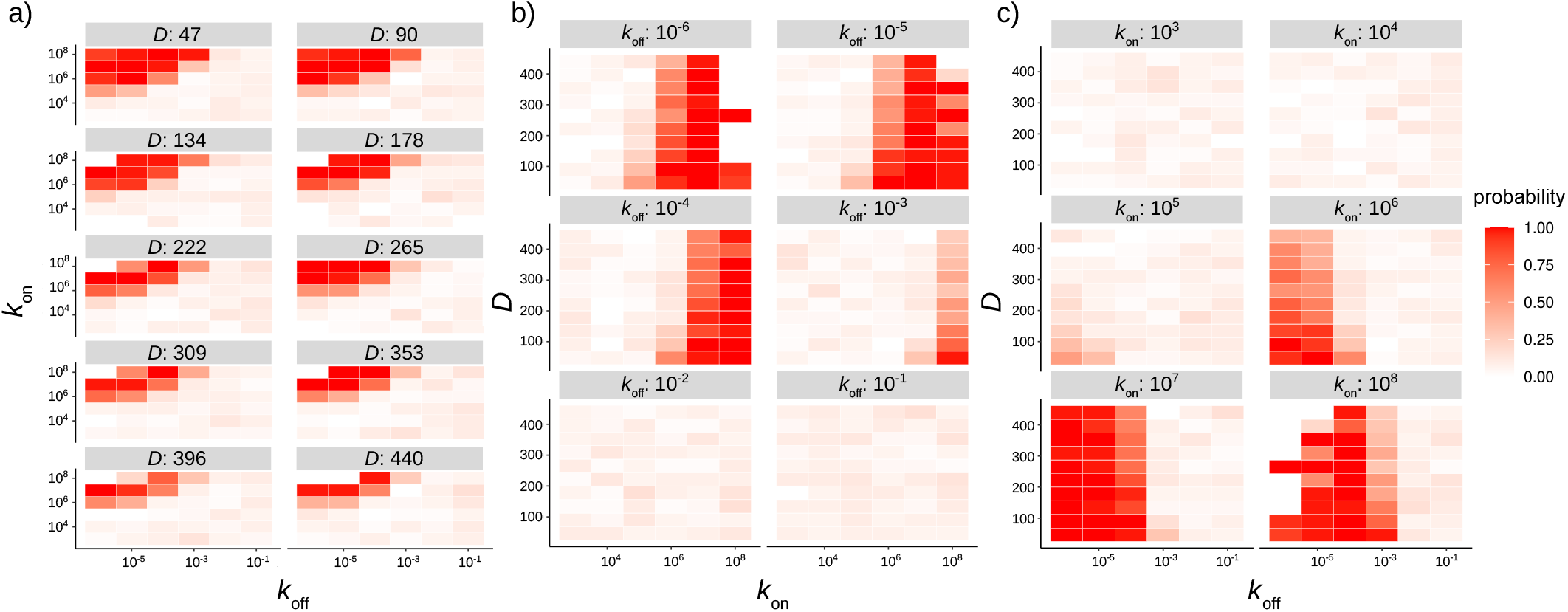
Establishment probability heatmaps for different two-dimensional parameter subspaces: a) *k*_off_ vs. *k*_on_ heatmaps for six values of *D*. b) *k*_on_ vs. *D* heatmaps for six values of *k*_off_. c) *k*_off_ vs. *D* heatmaps for six values of *k*_on_. Color denotes the magnitude of the probability, as indicated in the legend.

The conditions under which PG producers prevail can be intuitively understood as follows. The more bound molecules a PG cell has, the more fit it is, and thus more likely to establish. Lower diffusivity *D* slows PG diffusion, thus allowing PGs to spend more time in the producer clone vicinity leading to producers capturing a higher percentage of PGs. As the association constant *k*_on_ increases, producers capture more PGs per unit time, while increasing the dissociation constant *k*_off_ imparts the opposite effect.

Fig. 2a illustrates these trends well. A tug-of-war between *k*_on_ and *k*_off_ can be seen alongside the role of diffusivity *D*: lower diffusivity, in the top-left corner, maximizes invasion probability. *D* and *k*_off_ work in tandem, as seen in Fig. 2c, with high values of *D* and *k*_off_ associated with reduced invasion probability. We note that for low enough values of *k*_on_ or high enough values of *k*_off_, invasion is unlikely for all biologically relevant values of the other two variables, as can be seen from the bottom two heatmaps of Fig. 2b and top two heatmaps of Fig. 2c.

### A radially-symmetric, analytical Moran model of public good production and invasion

To further understand the model behavior and its generalizability, we complemented numerical simulations with an analytically-tractable modified Moran model of invasion probability. Here we summarize the main features of the approach (full derivation and details in Supplementary Information, SI). Fig. 3 illustrates the

**Figure 3.**
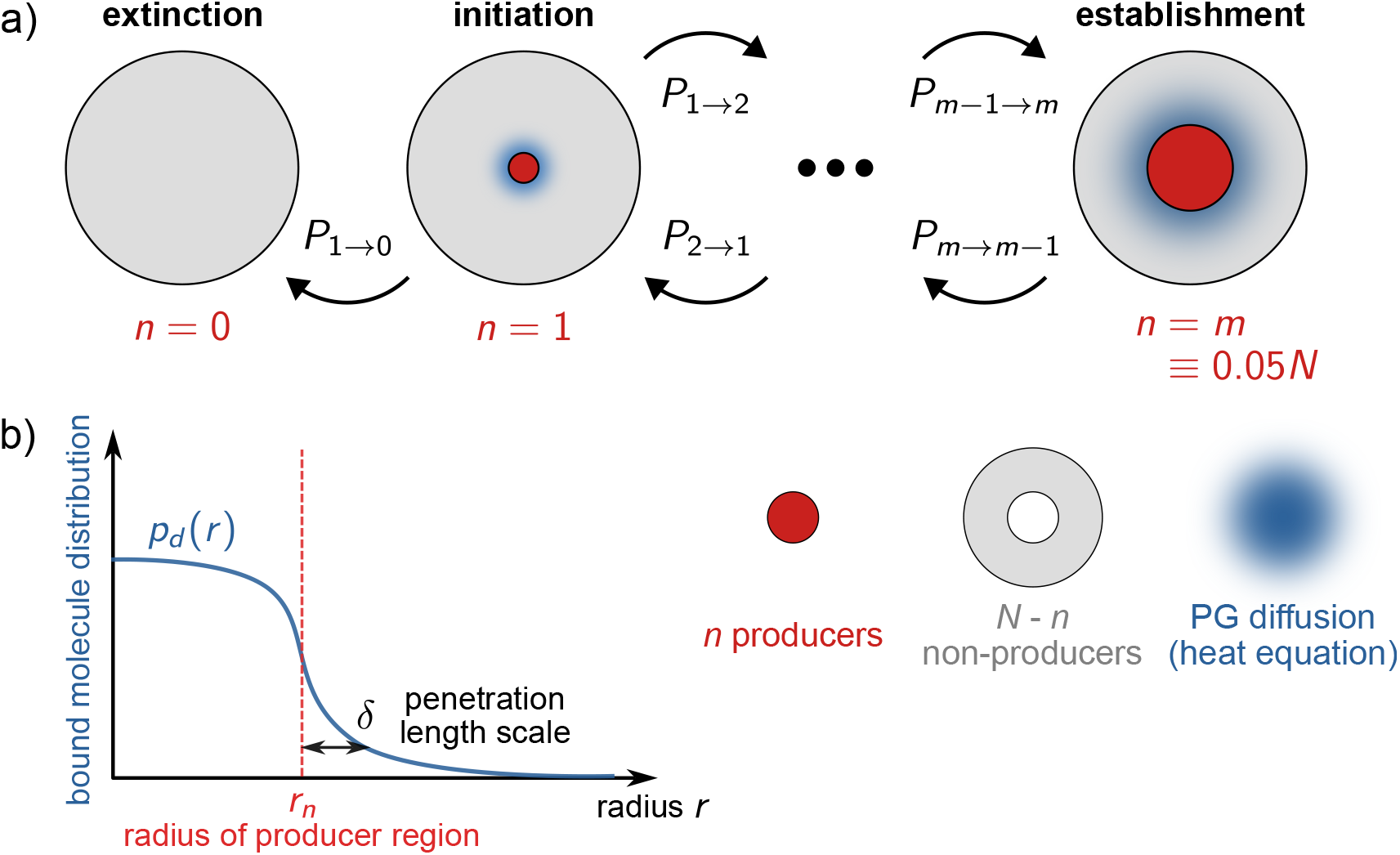
Overview of the analytical approach. *N* cells in *d* dimensions are arranged in a simplified radially symmetric geometry. There is a central producer region (red, consisting of *n* cells) surrounded by non-producers (gray, *N* − *n* cells). The dynamics are described with an effective Moran model, where the initial state has *n* = 0, and can transition via probabilities *P*_*n*→*n*+1_ and *P*_*n*→*n*−1_ to other states. Extinction of producers corresponds to reaching *n* = 0, while establishment is defined as reaching *n* = *m* = 0.05*N*. b) PG production leads to a steady-state radial probability density of bound PG molecules, *p*_*d*_ (*r*), that depends on the dimension *d* and the radius *r* from the center of the colony. Outside the producer region (which has radius *r*_*n*_ depending on the cell number *n*) the density falls off exponentially with a characteristic length scale *δ*.

Moran model consisting of a *d*-dimensional region of *n* producer cells at the center of *N* −*n* non-producer cells. For simplicity, the geometry is radially symmetric and the total number of cells *N* is fixed.

We investigate the probability *P*_est_ that producer cells (starting at *n* = 1) cross a threshold population size of establishment (defined as *n* = *m* ≡ 0.05*N*). This model is a Moran process because the *n* producers can stochastically increment by ±1, with the transition probabilities *P*_*n*→*n*−1_ and *P*_*n*→*n*+1_ controlled by the relative fitnesses of producers and non-producers. These fitnesses are, in turn, determined by the distribution of bound PG.

To estimate the distribution of bound PG, we use a continuous approximation of the reaction-diffusion model specified above—accounting for production, diffusion, binding/unbinding, and possible degradation or removal via processes like endocytosis. Simulations and time-scale analysis indicates that these processes are fast enough relative to the time scale of cell division that a steady-state can be reached, which allows us to obtain an analytical expression for the distribution *p*_*d*_ (*r*) of observing a bound molecule at radius *r* from the colony center. The distribution depends only on *r*_*n*_, the radius of the producer region, and a parameter 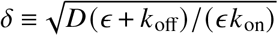, where *ϵ* is the degradation/removal rate. *δ* has units of length, and can be interpreted as a characteristic length scale determining how far *p*_*d*_ *r* extends into the non-producer region: *p*_*d*_ (*r*) ∼exp (−*r* /*δ*) for *r* ≫*r*_*n*_ (Fig. 3b). Smaller *δ* implies that more of the PG is localized to producers, giving a greater fitness advantage to the producer subclone — facilitating establishment.

This elegant formalism allows us to relate the establishment probability *P*_est_ directly to the characteristic length scale *δ*. In the Moran framework, *P*_est_ can be written in terms of the ratio *γ*_*n*_ = *P*_*n*_ →_*n*_ − _1_ /*P*_*n*_ →_*n*_ + _1_, the relative probability of the producers losing versus gaining one cell in the state *n*:

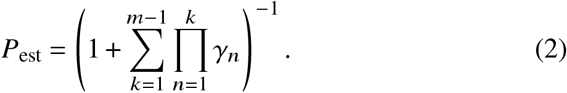

For our model the ratio can be well approximated by a simple expression,

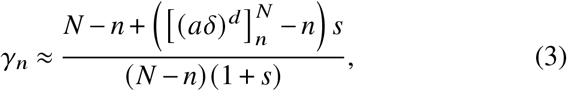

where *s >* 0 is a selection coefficient describing the maximum fitness benefit for a cell fully-saturated with bound PG molecules, and 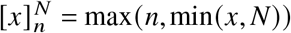 is a clamp function that restricts its argument *x* to be within the range *n* to *N*. The variable *a* sets an overall scale for the effect of *δ*, determined by the specific form of how the cell fitness depends on the bound molecule distribution.

Eqs. (2)-(3) reveal several insights into the underlying evolutionary dynamics. First, there is no explicit dependence of *P*_est_ on *k*_on_, *k*_off_, and *D*. These factors influence *P*_est_ only via *δ*. Specifically, what appears in Eq. (3) is *δ*^*d*^, which can be interpreted as a characteristic volume scale in *d* dimensions over which the public good is spread in the non-producer region. Hence, we predict that systems at a given *d* with the same value of *δ* should have the same *P*_est_ regardless of base parameter combination. This “curve collapse” of simulations is shown in Fig. 4a,b for *d* = 1 and *d* = 2, compared to the analytical model from Eqs. (2)-(3). The fitting parameters in the theory versus simulation comparison are *a* (the scale parameter which is fitted simultaneously to data from both *d* = 1 and *d* = 2) and the effective selection coefficients *s*, which are fit individually for each dimension. Second, while the stochasticity of PG production, diffusion, and (un)binding creates discernible deviations of simulations from continuous-approximation theories, overall agreement between the two approaches is quite good. Moreover the shift in *P*_est_ between *d* = 1 and *d* = 2 is also captured by the model.

**Figure 4.**
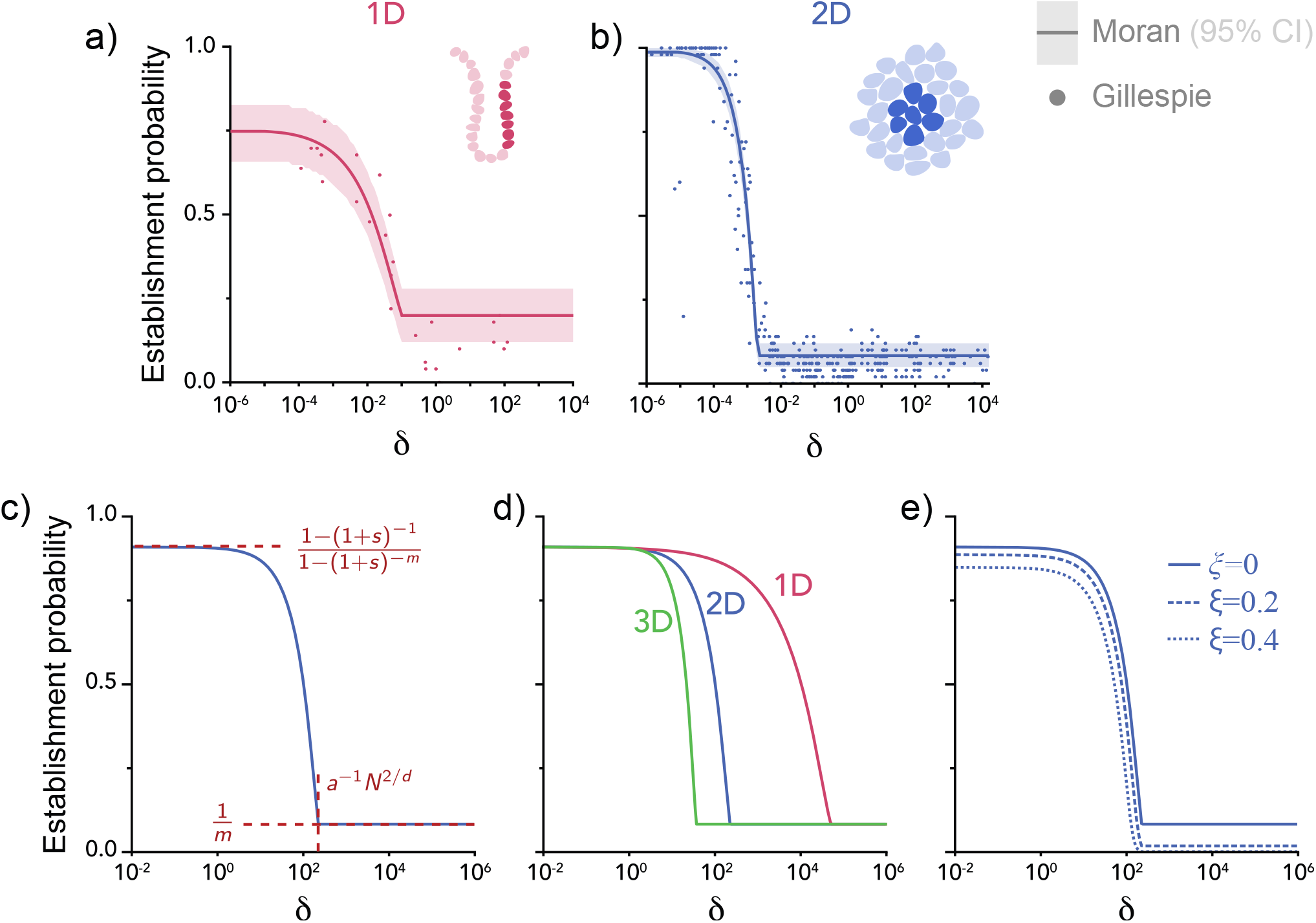
a,b) Numerical simulation (dots) and analytical theory (curves) results for the establishment probability in 1D and 2D systems, plotted as a function of the *δ* = *D* (*ϵ* +*k*_off_) /(*ϵ k*_on_). Though the simulations are run for a variety of different *D, k*_on_, and *k*_off_ combinations, they approximately collapse on a universal curve in terms of *δ*. This variable represents an effective length scale describing how far PG molecules penetrate into the non-producer region. c) An example establishment probability curve from the analytical theory, showing the two limiting values in the small and large *δ* regimes, as well as the threshold value of *δ* separting the regimes. d) Theoretical establishment probability curves in 1D, 2D, and 3D, showing the shift of the threshold to smaller *δ* with increasing dimension.e)The effects on establishment probability of the cost ξ, defined as the fraction of producer fitness lost due to PG production.

Eq. (3) also illustrates the limiting behaviors of invasion dynamics at large and small *δ*. For *δ* above a threshold value, *δ > δ*_*c*_ ≡*a*^−1^*N*^1/*d*^, PG is so diffusely spread out among producers and non-producers that each benefit equally (see Fig. 4c), and therefore establishment becomes effectively neutral, with *γ*_*n*_ = 1, and *P*_est_ = 1 / *m* — the neutral Moran model result for reaching an establishment population of *m* cells. For *δ < δ*_*c*_, PGs are largely confined to the producer region, and *P*_est_ increases with decreasing *δ*, reflecting the advantage gained by hoarding PG. In the limit *δ* →0, the public good does not spread beyond the producer cell, and is thus no longer “public”, and the producer clone establishes like an ordinary private mutation with *γ*_*n*_ = 1 / (1 + *s*)— the Moran model result for a mutant with a constant selection coefficient *s*. The resulting establishment probability in this limit is *P*_est_ = (1 − (1 + *s*) ^−1^) /(1 − (1 + *s*) ^−*m*^) (Fig. 4c). Overall, these limits agree well with the qualitative argument in Ref. 19 that the dynamics of public good traits depend critically on how well producers hoard public good.

The influence of tissue dimensionality is also described by Eq. (3). Larger *d* facilitates the spread of PG to non-producer regions, decreasing the threshold *δ*_*c*_ = *a*^−1^*N*^1/*d*^ for the neutral regime. We see this in Fig. 4d, where the threshold shifts to the left along the *δ* axis for theoretical *P*_est_ curves with increasing *d*. The dependence of *δ*_*c*_ on *N* can be also be qualitatively understood: larger *N* decreases the effects of genetic drift, reducing the probability that the mutant population goes extinct due to finite population sampling from generation to generation, thereby making establishment more likely for a given *δ*, increasing the threshold *δ*_*c*_.

Lastly, a fitness cost of producing public good can be incorporated into our model. We define *ξ* as the production costs normalized to maximum fitness gained by the PG, with 0 ≤ *ξ* ≤ 1. When *ξ* = 0 (nil production cost), we recoup our original Moran model. When *ξ >* 0, the formalism stays the same, except for a new ratio 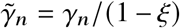 replacing *γ*_*n*_ in Eq. (2), where *γ*_*n*_ is given by Eq. (3). Fig. 4e depicts the effects of cost: increasing *ξ* creates an overall decrease in *P*_est_. The decrease is more pronounced at large *δ > δ*_*c*_, where the producer does not have preferential access to PG, but still pays the cost of production, making *P*_est_ significantly lower than the neutral Moran expectation. This high-diffusion limit confirms previous claims that that PG producers cannot establish in well-mixed populations^13^, and illustrates why it is necessary to carefully consider spatial structure when predicting the effects of ecological interactions on population dynamics.

## Discussion

Using both computational and analytical approaches, we found that public good phenotypes can invade and establish in cancer populations for realistic parameters when spatial structure and stochasticity are explicitly considered. The concordance between our two methodologies demonstrates that this conclusion is robust to modeling approach: Wright-Fisher vs. Moran proliferation, Cartesian vs. cylindrical coordinates, and on-lattice vs. continuous volume. Importantly, we identified a “super-variable” representing a characteristic length *δ*, that encapsulates variation in PG (1) association and (2) dissociation rates, (3) degradation/removal rate, and (4) diffusivity into a single descriptor of the PG mutation’s ability to selectively benefit producer cells and thereby invade into an evolving population. Curve-collapse in this variable shows that we have summarized the essential behavior of this system. By distilling model behavior into a single algebraic variable, we provide an intuitive understanding of when PG phenotypes can emerge in evolving tumors and when they cannot.

Stochasticity was essential to producer establishment in both models. To the best of our knowledge this is the first investigation of PG producer invasion into an existing tumor that directly incorporates stochasticity and spatial structure. Because mean-effect models cannot explain establishment of PG phenotypes in a tumor, we modeled every significant source of randomness that might reasonably explain how producer phenotypes establish in real tumors: diffusion, secretion/binding, and cell proliferation.

Producer clones establish in 1D, 2D, and 3D tissues. Invasions in 1D and 2D follow the same general shape although producer clones in 1D invade at higher values of *δ* (because 1D producers sequester a larger fraction of their PG) but plateau at a lower maximum invasion probability (because bound PG saturates at a lower level). We identified an analytical function describing establishment probability across dimensions, which agreed well with simulated 1D and 2D tumors. 1D tissue models are generally used to model intestinal stem cells compartments important in colon cancer)^29^, while 2D and 3D formalisms are generally used for stratified epithelial sheets and soft tissues^30^. Thus, 1D, 2D, and 3D models can in principle be experimentally validated.

Our study has limitations. While spatial evolution is generally modeled with either on-lattice or radially-symmetric models like ours, real tumor tissue is structurally heterogeneous and irregular. Additionally, our modeling requires knowing precise values of the biological parameters identified in Table 1 for any particular tumor and PG phenotype. Thus, while we studied a very broad range of biologically relevant parameters, only sub-ranges of this space might be applicable to each particular tumor/PG combination. Finally, we studied tumors at carrying capacity; expanding tumors might establish producer phenotypes differently. Along with addressing these outstanding questions regarding approach, it will be interesting to connect our low cell number stochastic regime to the previously investigated large cell number regime^9–12, 23^. In addition, running numerical simulations in three dimensions and more complex geometries, along with including a more comprehensive model of fitness costs associated with PG production, will be valuable extensions of the study.

While our analyses focused on explaining a paradox in cancer clonal dynamics, we believe that our mathematical formalism may have utility in prognosis and treatment — most therapeutic agents mimic public goods. Our model of subclone competition has immediate relevance to therapeutic resistance and current clinical approaches to mitigate it that try to eliminate resistant subclones using non-resistant competitors^31–35^. Moreover, other diffusible molecules that play a role in treatment could be added to the model, for example molecular chemotherapies that stochastically regulate cancer cell division and death, e.g. growth or cytokine inhibitors, or angiogenic/hypoxia regulators.

## Methods

### Numerical model

Stochastic spatial simulations were performed in Python 3.8.10 and 3.10.8. A two-dimensional 15 × 15 square lattice with a von Neumann neighborhood was used, with nodes representing tumor cells. While 2D lattices do not fully recapitulate 3D tissue structure^36^, they have a long history of use in modeling PGs in tumor evolution^9–12, 14^ and are noted to be a good representation of a simple epithelium at early stages of carcinogenesis^14^. Cells are represented as point-like objects, each having 25,000 receptors^37^, with separation of Δ*x* = 14.6 *µ*m calculated from tumor cell density given in Ref. 38.

Public good (PG) factors were secreted every 1.95 seconds (value for VEGF in adipocyte cells calculated from the data in Ref. 39). 1,000 molecules were secreted each time to accelerate simulations. PG factor diffusion was modeled as a random walk on the lattice, with each step occurring in only one of the two possible dimensions (dubbed as the ‘OR move’ in Ref. 40). Ten values of diffusivity (see Table 1) were used covering the full biologically relevant range of measured diffusivities^28^.

Each diffusion time step is approximated as a time interval Δ*t*_diff_ = MSD / (4*D*), where *D* is the diffusivity^41^ and the mean square displacement MSD is taken to be 1 *µ*m^2^ for simplicity. During this step, diffusing molecules have a chance to bind to the closest cell represented by a lattice node and ligands already bound to cells have a chance to escape. The stochastic association-dissociation process for each cell was implemented following the classical Gillespie algorithm^42^, with the on-rate normalized by the volume and Avogadro’s number as described in Ref. 37. The details of the algorithm are as follows:

1. Initialize current step time: *t*_step_ = 0
2. Calculate auxiliary variables:
  a. *a*_1_ = *k*_on,norm_ × (# free molecules around cell) × (# receptors per cell), where *k*_on,norm_ is the normalized on-rate, *k*_on,norm_ = *k*_on_ /(*N*_*a*_*V*_node_), *k*_on_ being the raw on-rate, *N*_*a*_ being the Avogadro number and *V*_node_ the volume of the cell, *V*_node_ = Δ*x*^3^.
  b. *a*_2_ = *k*_off_ × (# molecules bound to cell), where *k*_off_ is the off-rate
  c. *a*_0_ = *a*_1_ + *a*_2_
3. Generate two random numbers *ρ*_0_ and *ρ*_1_, each uniformly distributed between 0 and 1.
4. Calculate time till the next reaction: 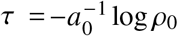.
5. Recalculate current step time: *t*_step,new_ = *t*_step_ + *τ*. If *t*_step,new_ *>* Δ*t*_diff_, no reaction happens at this cell during the current diffusion step, otherwise decide which reaction, association or dissociation, will take place: if *ρ*_1_*a*_0_ ≤ *a*_1_ and (# free molecules around cell)*>* 0 or if (# molecules bound to cell) = 0^1^, free molecule-cell association occurs, otherwise bound molecule-cell dissociation occurs.
6. Items 2 – 5 are repeated until *t*_step,new_ *>* Δ*t*_diff_.

Six values of association and dissociation constants covering the full biologically relevant range of measured values were used (Table 1).

The diffusion/binding procedure was repeated multiple times until the time of proliferation (taken as 24 hours^43^) arrived. Then the bound molecules were mapped to cell “benefits” using the Hill function in Eq. (1) with *n* = 1. The constant EC_50,bound_ is the usual half-maximal effective concentration EC_50_ converted to the number of molecules: EC_50,bound_ = EC_50_ *N*_*a*_*V*_node_ and *N*_bound,cell_ is the number of molecules bound to the cell. An EC_50_ value of 7.76 × 10^−11^ M for VEGF165-VEGFR2 binding from Ref. 44 was used.

The proliferation step was implemented using Wright-Fisher procedure^23^ (see Figure 1): each cell contributed a number of putative offspring proportional to the cell fitness to a common pool (a coefficient of proportionality of 1,000 was used). The pool was then randomly sampled. This regular Wright-Fisher procedure was complemented by enforcing the clustering of those sampled offspring that were producers at the center of the lattice to keep the proper spatial structure.

The proliferation/diffusion/binding procedure was repeated until the fraction of producers reached 0.05. The invasion threshold of 0.5 was also tried, but it was found that the difference with the threshold of 0.05 consisted mostly in the computational resources used. The simulation was run until the threshold was reached (at which point the producer invasion was considered as having occurred) or until there were no producers left (producer perishing has occurred).

The one-dimensional simulation was performed in the same fashion, only the two-dimensional lattice was substituted by a 100-cell long one-dimensional line and the initially invaded producer was put in the center of the line.

## Analytical model

Full details of the analytical Moran model are described in the SI.

## Supplementary Information

### 1 Moran model derivation

#### 1.1 Geometry of the cellular population

We consider the cellular colony as a total population of *N* cells filling up a *d*-dimensional region of size *S*_*d*_ (*r* _*N*_) characterized by radius *r* _*N*_. For *d* = 1 this region is a line of length *S*_1_(*r* _*N*_) = 2*r* _*N*_, for *d* = 2 a circle of area 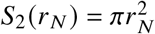 and for *d* = 3 a sphere of volume 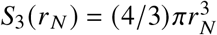. For *d* ≥ 2 we will adopt radial coordinates, with *r* = 0 corresponding to the center of the region (the origin) and *r* = *r* _*N*_ the edge of the colony. For *d* = 1 we have a line segment along the *r* axis: −*r* _*N*_ *< r < r* _*N*_. We define *r*_1_ as the effective radius of a single cell, such that *S*_*d*_ (*r*_1_) = *S*_*d*_ (*r* _*N*_) / *N* is the size of the *d*-dimensional region containing an individual cell. Assuming *r*_1_ is fixed, we can solve for *r* _*N*_ in terms of *r*_1_ and *N*: *r* _*N*_ = *r*_1_*N*^1/*d*^. At any given time there are *n* producer cells and *N* −*n* non-producer cells. Assuming the producers were seeded at the center of the region, they form a subcolony of size *S*_*d*_ (*r*_*n*_) at the origin, where *r*_*n*_ = *r*_1_*n*^1/*d*^.

#### 1.2 Production and diffusion of the public good

We assume that production, diffusion and binding of the public good molecule is sufficiently fast that the concentrations of molecules in solution reaches a steady state very quickly on the timescale of a single Moran evolutionary step (i.e. the time scale of cell division). We define *u* (*r*) as the steady-state *d*-dimensional density of free molecules (with units of length^−*d*^). Note that by symmetry this density only depends on the radial distance *r*. For *d* = 3 the density can be interpreted as concentration (units of inverse volume). When *d <* 3 we can convert it to a concentration by assuming cross-sectional dimensions for our colony. For example if *h* is the perpendicular thickness of the circular colony in *d* = 2, then its total volume is *S*_2_(*r* _*N*_)*h* and *u*(*r*)/*h* is a concentration. Similarly if *h*^2^ is the cross-sectional area of the *d* = 1 linear colony, its total volume is *S*_1_(*r* _*N*_)*h*^2^ and *u*(*r*)/*h*^2^ is a concentration.

In the steady state the free molecule density *u*(*r*) satisfies the following reaction-diffusion equation:

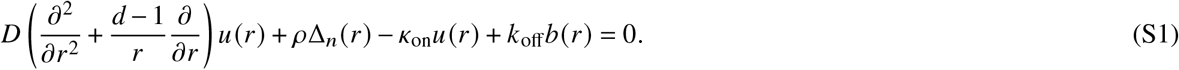

The first term of Eq. (S1) describes diffusion of the free molecules with diffusivity *D*, where the operator in parentheses is the radial component of the *d*-dimensional Laplacian. The second term of Eq. (S1) describes the production of public good with rate *ρ* per producer cell (units of inverse time). Δ_*n*_ (*r*)encodes the spatial distribution of the production when there are *n* producer cells: Δ_*n*_ (*r*) = 1 /*S*_*d*_ (*r*_1_) for *r* ≤ *r*_*n*_ (within the producer region); Δ_*n*_ (*r*) = 0 for *r > r*_*n*_ (outside the producer region). The form of Δ_*n*_ (*r*) means that total production of public good is just *n* times the single-cell rate *ρ*:

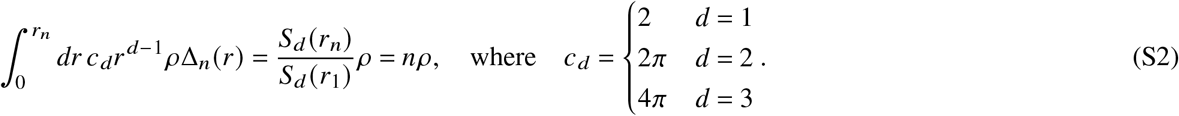

Here *c*_*d*_ is the solid angle coefficient in *d*-dimensions, ensuring the correct volume element in the integration: 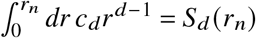

The third and fourth terms of Eq. (S1) describe respectively the loss of free molecules due to binding, −*κ*_on_*u*(*r*), and the gain of molecules from unbinding, *k*_off_*b*(*r*), where *b*(*r*) is the density of bound molecules. In this framework *κ*_on_ has units of inverse time. To relate *κ*_on_ to a biochemical binding constant *k*_on_ (with typical units of M^−1^s^−1^), we need to know the concentration [*R*] of receptors in any given region of our colony (which itself is a product of the number of receptors per cell times the concentration of cells). Then *κ*_on_ = *k*_on_ [*R*]. Note that *κ*_on_ has the same value even when *r > r* _*N*_, implicitly assuming that there are other cells (or the surrounding medium) which absorb molecules at large distances. However since we typically work with *N* ≫ *n* and hence large *r* _*N*_, the density *u* (*r*) is usually vanishingly small at *r* _*N*_, so the details of the region beyond are irrelevant to the results.

The density of bound molecules *b* (*r*)can be related to *u* (*r*) by assuming a steady state between binding, unbinding, and degradation (or removal via endocytosis) of molecules:

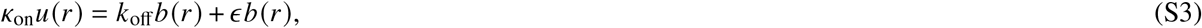

where *ϵ* is the degradation rate. In the simulations this degradation happens during reproduction at the end of each generation (once per 24 hours). Hence we set *ϵ* = (24 hr)^−1^ = 1.16 × 10^−5^ s^−1^ to match the simulation timescale. Solving for the steady state *b*(*r*) from Eq. (S3) and plugging it into Eq. (S1), we can express the latter entirely in terms of *u*(*r*),

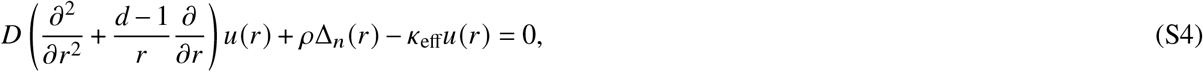

where we introduce an effective rate:

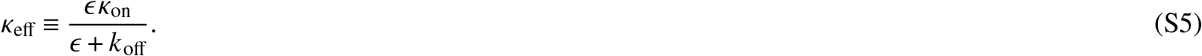

Integrating Eq. (S4) over all space, and using Eq. (S2), we get the following relation for the integral of the steady-state density of free molecules:

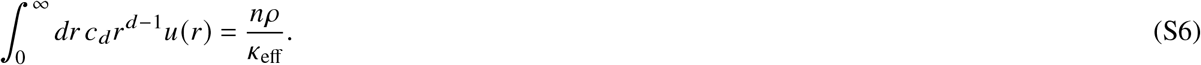

Since we know from Eq. (S3) that the bound density *b* (*r*) = (*κ*_eff_ /*ϵ*) *u* (*r*), we can equivalently express Eq. (S6) in terms of the bound density as:

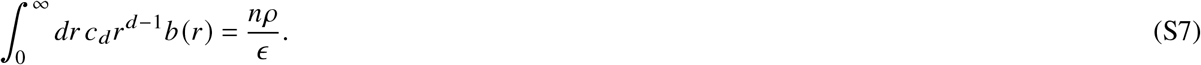

Equation Eq. (S4) is not analytically solvable for *u* (*r*), but there is an approach that yields an approximate closed form expression in all dimensions. If we replace the production term Δ_*n*_ (*r*) by an equivalent Dirac delta source concentrated at the origin of magnitude *nρ*, the equation can then be solved, and the density profile still satisfies Eq. (S6). The approximate *u* (*r*) agrees with the true *u* (*r*) at *r* ≫ *r*_*n*_, away from the source, but diverges as *r* → 0. To get a better overall approximation to the real profile (removing the divergence), we thus replace *u* (*r*) for *r < r*_*n*_ by a constant, chosen to ensure Eq. (S6) still holds. This roughly reflects the saturation in concentration that occurs within the producer region. Using Eqs. (S6)-(S7), we can express the resulting answers in normalized form, as *p*_*d*_ (*r*) = *u*_*d*_ (*r*)*κ*_eff_/(*nρ*) = *b*_*d*_ (*r*)*ϵ* /(*nρ*), where the subscript *d* refers to the dimension:

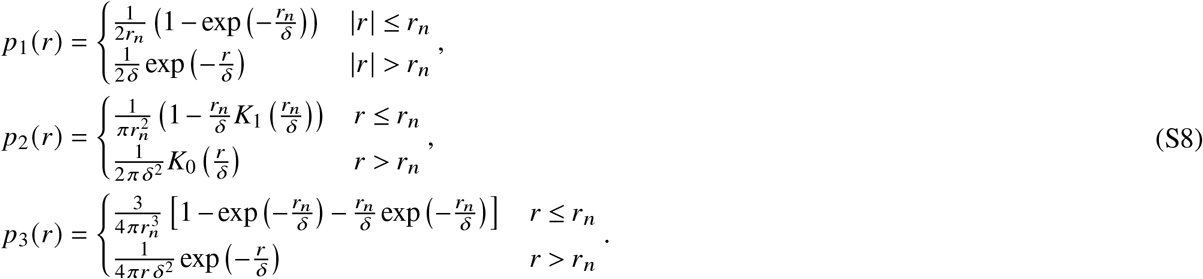

Here 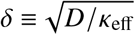, and in the expression for *p*_2_ (*r*) the functions *K*_*v*_ (*x*) for *v* = 0, 1 are modified Bessel functions of the second kind. *p*_*d*_ *r* can be interpreted as the probability density in the steady state of observing a molecule at *r*, conditional on it being bound (or equivalently conditional on it being unbound). It satisfies the normalization 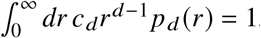. Note that from Eq. (S8) we see that *p*_*d*_ (*r*) depends only on two parameters, *r*_*n*_ (the radius of the producer region) and *δ*. The latter has units of length, and can be interpreted as the effective distance over which molecules diffuse beyond the producer region. For *δ* → 0 we see that *p*_*d*_ (*r*) vanishes for *r > r*_*n*_ (no molecules outside of the producer region), while for *δ* → ∞ the probability density becomes uniform across the entire space.

#### 1.3 Moran model

We assume the fitness of a cell depends on the presence of bound molecules, represented by the probability density *p*_*d*_ (*r*). We thus take a simple exponential sigmoidal form to capture this dependence, with the fitness *f*_*d*_ (*r*) defined as:

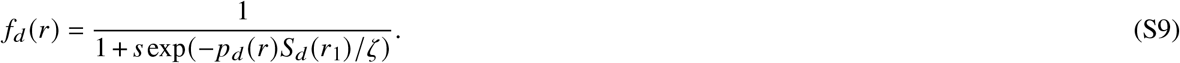

In the exponential expression we use *p*_*d*_ (*r*)*S*_*d*_ (*r*_1_) to represent the conditional probability of finding a bound molecule in the volume corresponding to a single cell. For low probabilities *p*_*d*_ (*r*)*S*_*d*_ (*r*_1_) ≪ *ζ* below some threshold *ζ*, we have *f*_*d*_ (*r*) ≈ 1/(1 + *s*), and for high probabilities *p*_*d*_ (*r*)*S*_*d*_ (*r*_1_) ≫ *ζ* we have *f*_*d*_ (*r*) ≈ 1. Here *s >* 0 is the selection coefficient describing the fitness benefit from having bound molecules. Then mean fitness 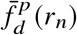 of a cell in the producer region is given by integrating the fitness *f*_*d*_ (*r*) over the entire region and dividing by the area:

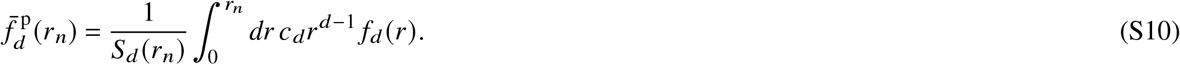

Similarly the mean fitness of a non-producer cell 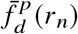 can be expressed as:

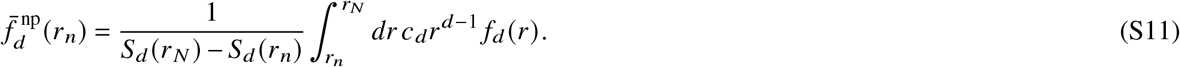

To make the connection with a Moran model, we note that in an ordinary two-allele Moran model (*n* mutants, *N* − *n* wild types) the key quantity is *γ*_*n*_ = *P*_*n*→*n*−1_/*P*_*n*→*n*+1_. This is the ratio of the transition probability *P*_*n*→*n*−1_ to lose a mutant in the next time step and the probability *P*_*n*→*n*+1_ to gain a mutant. It is related to the fitness via 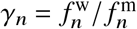, where 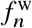 is the fitness of the wild type and 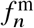 the fitness of the mutant (which can depend on the current state of the population described by *n*). In our case we replace mutants by producers, and wild types by non-producers, and define 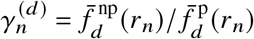. We can now use all the standard results from the theory of Moran processes. In particular, the establishment probability *P*_*d*_ (*m*), that a single producer introduced into the system will reach a population size *m*, is given by:

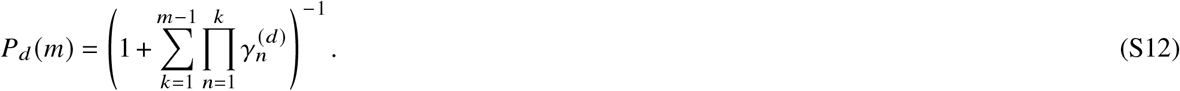

This probability depends on *δ*, and reduces to conventional Moran results in both the *δ* → 0 and *δ* → ∞ limits. In the first limit, *δ* → 0, the probability density of public good for non-producers vanishes, *p*_*d*_ (*r*) = 0 for *r > r*_*n*_. Hence only producers get a fitness benefit, with 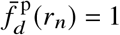 and 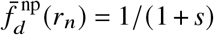. We thus get 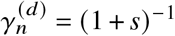 and the establishment probability becomes

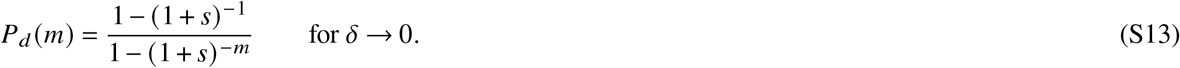

In the second limit, *δ* → ∞, the public good is spread out evenly to producers and non-producers, with *p*_*d*_ (*r*) approaching a constant throughout the entire region. Thus 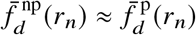 and 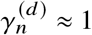. The establishment probability is the neutral Moran result, which is the *s* → 0 limit of Eq. (S13),

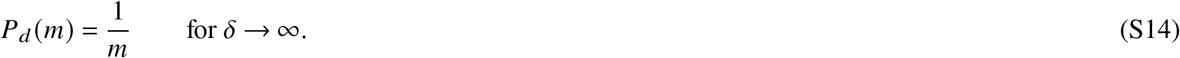

For intermediate *δ* the probability *P*_*d*_ (*m*) is a sigmoidal function that decreases from the value of Eq. (S13) to that of Eq. (S14). It is possible to closely approximate this sigmoid using an analytical expression for *γ*_*n*_. To derive this approximation, we focus on the regime where the threshold value *ζ ≪* 1. Since *p*_*d*_ (*r*) *S*_*d*_ (*r*_1_) is a conditional probability for a single molecule to be bound to a cell, and there are a large number of public good molecules produced, even for small values of *p*_*d*_ (*r*) *S*_*d*_ (*r*_1_) there are likely to be multiple bound molecules overall. We can think of *ζ* as roughly describing the relationship between the magnitude of public good production and the associated fitness effects: smaller *ζ* corresponds to more molecules produced, since one sees the same fitness effects (proportional to total number of bound molecules) for a smaller values of the single molecule bound probability *p*_*d*_ (*r*) *S*_*d*_ (*r*_1_). We assume *ζ* ≪ 1 (large production) as the biologically relevant regime. In this case one can show that the integrals in Eqs. (S10)-(S11) can be simplified, so that their ratio is approximately given by:

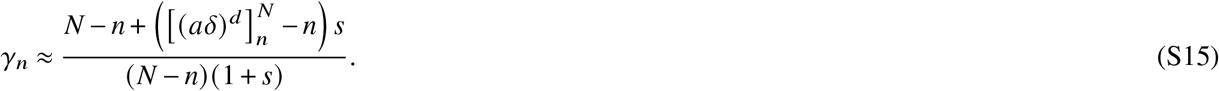

Here 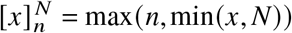 is a clamp function that bounds its argument *x* to be within the range *n* to *N*. The variable *a* ≡ log *ζ* essentially sets an overall scale for *δ*, shifting the entire sigmoid on the *δ* axis. In comparing to simulation data we just use this scale as an extra fitting parameter.

To incorporate the effect of public good production coming with a metabolic cost for the producer, we can use a simple modification, replacing 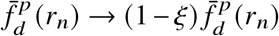, where 0 ≤ *ξ* ≤ 1 reflects the percentage of producer fitness lost due to costs. This changes Eq. (S15) to:

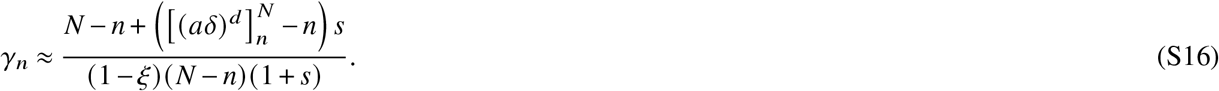

## Footnotes

1 If (# free molecules around cell)= 0 and (# molecules bound to cell)= 0 the cell is skipped altogether.

## Notes

### Competing Interest Statement

The authors have declared no competing interest.

